# An Integrated Approach to Late Chalcolithic Bone Artefacts from Yeghegis-1 (Armenia): Technological, Traceological, and ZooMS Analyses

**DOI:** 10.1101/2025.08.27.672598

**Authors:** Évora Marina, Mkrtchyan Satenik, Saribekyan Mariam, Amano Noel, Thuering Ulrrike, Schimer Shtefanie, Yepiskoposyan Levon, Roberts Patrick, Antonosyan Mariya

## Abstract

This article presents a preliminary study of the osseous industry from the Yeghegis-1 rock shelter in southern Armenia, dated to the Late Chalcolithic period. An integrated methodological approach, combining technological analysis, typological classification, use-wear studies, and Zooarchaeology by Mass Spectrometry (ZooMS), is applied to the assemblage. Here we reconstruct production strategies, use patterns, and discard behaviours associated with osseous material processing. Initial results reveal a preference towards the use of easily accessible Caprine bones for bone manufacture. Reconstructed technological patterns are broadly consistent with other regional Chalcolithic traditions, while also pointing to distinct local practices. This study contributes to a broader understanding of prehistoric technological systems in the Lesser Caucasus, highlighting the role of osseous industries in the daily lives of Chalcolithic communities.

## 1. Introduction

The analysis of osseous industries offers critical insights into the technological, economic, and social dynamics of prehistoric communities. Osseous raw materials like antlers, mammal bones, and ivory played a fundamental role in tool production, complementing lithic and wood technologies and serving a wide range of utilitarian and symbolic functions (Bicho et al., 2004; Évora, 2013). Despite the increasing recognition of their importance in prehistoric contexts, osseous tools remain understudied in archaeological research in the Lesser Caucasus. Research has traditionally focused on typological and technological aspects, with little focus on understanding the selection of source materials and the role animals played in daily lives of prehistoric communities. However, recent methodological advancements, particularly the application of palaeoproteomics, allow the identification of the animal species used in bone tool manufacture. One such method, Zooarchaeology by Mass Spectrometry, was applied to a curated museum assemblage from Armenia for the first time in the South Caucasus region, marking a significant advancement in the study of prehistoric bone tool manufacture in the region (Antonosyan et al., 2025a).

The earliest known presence of osseous artefacts in the Caucasus dates back to the late Middle - Upper Palaeolithic, mainly represented by bone points, awls, hunting weapons, eye needles, and personal ornaments (Yeritsyan 1982; Adler et al. 2008; Golovanova et al., 2010; Bar-Yosef et al., 201; Kandel et al., 2017). The intensity, diversity, and functionality of tool production increased in the Pottery Neolithic (6000-5200 BCE). With the emergence of agriculture in the Neolithic, the intensity, diversity, and functionality of bone manufacturing increased. The regional Neolithic bone artefact composition includes personal ornaments and implements for farming, pottery making, textile or leather making, wood or stone processing, gaming, etc (Sagona 2017; Zhvania 2017; Taha 2017; Hayrapetyan et al., 2018; Arai et al., 2021; Eloshvili 2021; Chataigner et al., 2022). In the Chalcolithic (5200-3500 BCE), significant socio-economic transformations occurred, including shifts in subsistence strategies, settlement patterns, and material culture. In this context, the study of bone tool production can shed light on broader aspects of technological organisation, specialisation, and adaptation.

In the Chalcolithic of Armenia, bone tools related to domestic production are reported in several sites (Choyke, 2000; Chataigner et al., 2010; Stapleton et al., 2018; Zarikian and Kalantaryan, 2021); however, compelling evidence for specialised bone tool production is reported in two key sites: Areni-1 cave and Getahovit-2 cave. These sites, within the broader technological and cultural landscape of the Lesser Caucasus, reveal distinct patterns of osseous tool manufacturing and function. At Areni-1 cave (4300-3500 cal. BCE, Areshian et al., 2012; Wilkinson et al., 2012), the excavations revealed an exceptional preservation of organic materials and tools used in both subsistence and ritual contexts (Areshian et al., 2012; Wilkinson et al., 2012; Stapleton et al., 2018; Samei et al., 2020). This dual functionality underscores their integration in daily life and symbolic practices, like the bone awls and needles found in funerary contexts (Samei et al., 2020). At Getahovit-2 cave (ca 5200-4900 cal BCE, Kakantaryan and Ghanem, 2019), the bone tool industry, alongside the lithic industry, offers insights into Chalcolithic pastoral economies and their technological adaptations (Kakantaryan and Ghanem, 2019; Zarikian and Kalantaryan, 2021). The cave served as a seasonal shelter for shepherds, with an osseous industry manufactured *in situ*, utilising locally available Caprine and cervid remains (Zarikian and Kalantaryan, 2021), for tasks such as hide-working and textile production. Here, tools like scrapers, perforated ribs, and needles were reported, similar to some pastoralist communities in Anatolia, such as Uğurlu (Paul and Erdoğu, 2017; Kalantaryan and Ghanem, 2019; Kalantaryan et al., 2022). However, the osseous tools technological and use-wear research is more common in some European and other Middle Eastern regions (Choyke, 2000; Russel, 2001; Choyke and Bartosiewicz, 2004; Sidéra, 2006; Crabtree and Campana, 2010; Vitezovic, 2021; Mărgărit et al., 2022, 2023; Skakun et al., 2025).

Among the Armenian archaeological sites cited above, Yeghegis-1 rock shelter (Figure 1) provides an excellent opportunity to examine the Chalcolithic bone tool assemblage recovered from a stratified and well-documented context. Excavated in recent years, Yeghegis-1 has yielded a diverse range of osseous artefacts, reflecting complex technological choices and craft traditions. This paper presents a preliminary analysis of the osseous industry from Yeghegis-1, focusing on five main dimensions: 1) analysis of the technological traits, such as manufacturing techniques, and possible evidence of standardization or variability; 2) identification of functional attributes through macroscopic and microscopic use-wear analysis; 3) typological classification of the bone tool artefacts according to form, function, and morphological features; 4) species identification of animal remains employed in bone manufacturing using minimally destructive palaeoproteomics technique; and 5) contextualization the assemblage within the broader framework of Chalcolithic osseous industries in the region. Our goal is to contribute to the growing body of knowledge on Chalcolithic technologies and human-animal interactions in the Lesser Caucasus and to position the Yeghegis-1 assemblage within regional framework of osseous tool production and use.

**Figure 1.**
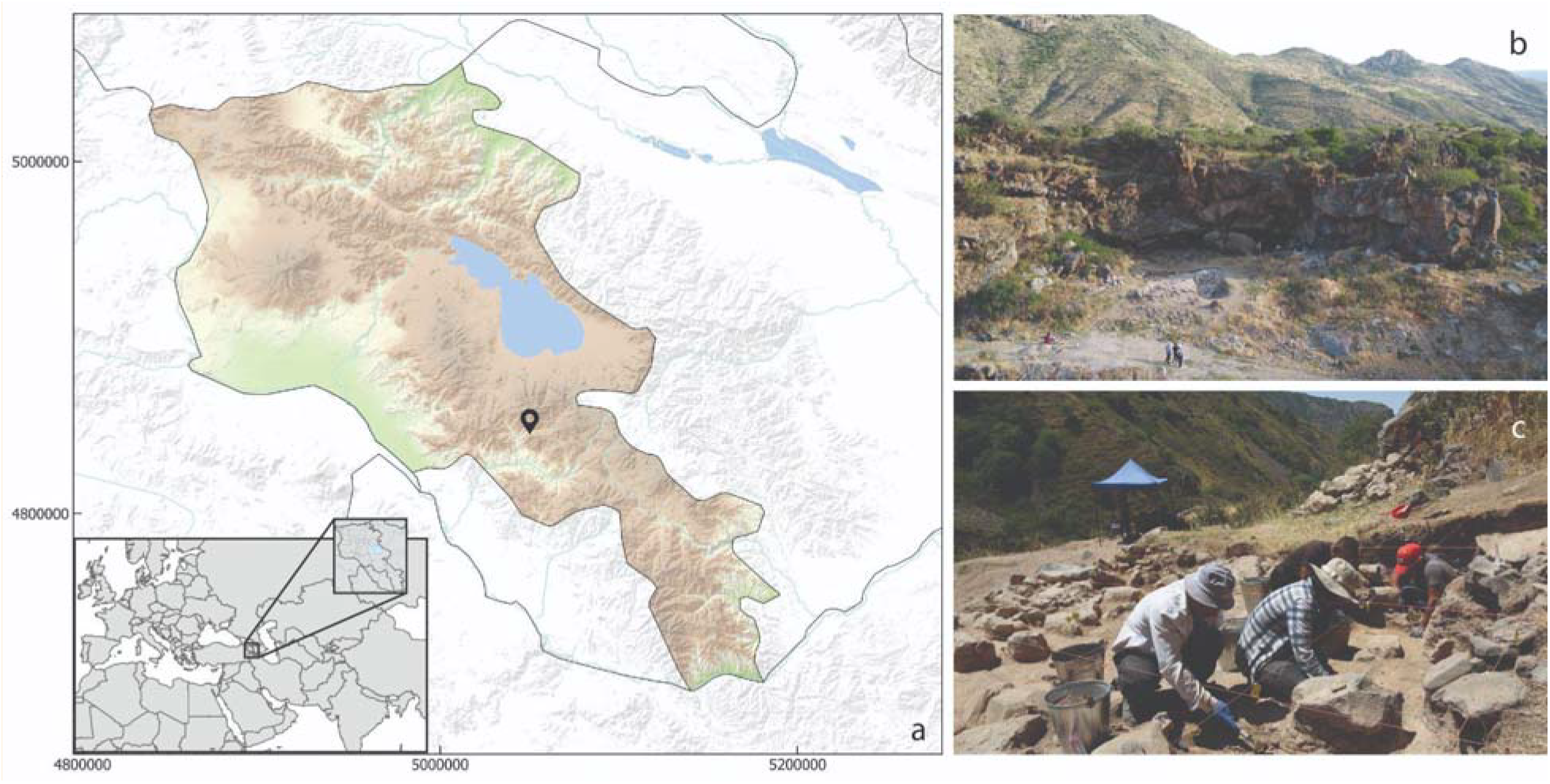
a) Yeghegis-1 archaeological site location in Armenia; b) general view of the archaeological site; c) partial view of the excavation during the 2024 fieldwork.

## 2. Site description

Yeghegis-1 is a basalt rock shelter situated in the Yeghegis River valley, at an elevation of 1500 m above sea level in Vayots Dzor Province, Armenia. The excavations at the site yielded a stratified sequence of five artefact-rich (ceramics, lithics, copper ores, charcoal, and animal bones) occupation layers, called Horizons. Radiocarbon dating of these layers confirms 600 years of human occupation at the site spanning from ca. 4100 cal BCE (Horizon 5, the lowest excavated layer) to ca. 3500 cal BCE (Horizon 0, top layer) (Frahm et al., 2024; Antonosyan et al., 2024; 2025b), situating the site within the Late Chalcolithic period of the South Caucasus (4300–3500 BCE; Bobokhyan et al., 2014; Sagona, 2017).

The lithic assemblage at the site is predominantly composed of obsidian artefacts, despite the absence of local obsidian sources in the river valley. Geochemical sourcing of obsidian artefacts indicated that communities utilised a diverse array of raw material sources situated at varying distances from the rock shelter, ranging from approximately 40–55 km to as far as 220 km in linear distance. At the same time, the analysis revealed an increase in obsidian diversity through time, suggesting shifts in land use and, in turn, social connections across the wider region (see Frahm et al., 2024, for the detailed description of the lithic assemblage).

Archaeobotanical analysis at the site revealed a small assemblage of cultivated crop remains represented by barley and wheat, accompanied by pulses represented by lentils, cotyledons, and chickpeas. The macrobotanical assemblage also includes wild (presumably foraged) berries, fruit (n=107), and nut trees, all of which grow wild around the site and are native to the region (see details in Antonosyan et al., 2025b). While the present data are too limited to reconstruct the role of cultivation in the economy at the site, the absence of agricultural tools and plant chaff may suggest a low-investment form of cultivation and opportunistic collection of wild plants.

The faunal assemblage is dominated by Caprines (both sheep and goat were identified via ZooMS technique) throughout the period of site occupation, highlighting the importance of these medium-sized ungulates in the subsistence economy of the groups that occupied the site. The second dominant group is cattle, less frequent than Caprines, reflecting their limited importance in the animal economy. Wild fauna is represented by gazelles, boars, hares, birds, and fish. A broad range of carnivore taxa was also recorded, including mustelids, wolves, foxes, wild cats, and bears (Antonosyan et al., 2025b). The presence of wild mammals, along with fish and bird remains, indicates a broader exploitation of natural resources, complementing pastoralism. However, their low frequency suggests that this was likely opportunistic rather than a primary subsistence strategy. Herd mobility and management strategies at the site were assessed through stable oxygen, carbon, and nitrogen isotope data that suggest a settled pastoral system with year-round reliance on local highland pastures (see details in Antonosyan et al., 2025b).

## 3. Materials and Methods

### 3.1 Sampling

The first brief scientific description of the site was completed in 2020, followed by test excavations in 2021. Large-scale excavations at the site started in 2022 and continued in 2023 and 2024. A trench measuring 6 x 4 m was excavated in square meter units to a depth of 2 m. All materials were collected and recorded according to the established stratigraphic divisions (Horizons and Subhorizons). Bones and other fossils recovered during the excavation were collected *in situ*, and their stratigraphic position was recorded. Excavated sediments were removed for dry sieving with 2- and 0.5-mm sieves to recover smaller specimens. All finds were brushed and cleaned in a field laboratory and stored in airtight, opaque bags. The stratigraphic sequence of the Trench was divided into six Horizons (H0, H1, H2, H3, H4, H5), further subdivided into Subhorizons (H1Sp1, H1Sp2, H2Sp1, H2Sp2, H3Sp1, H3Sp2, H4Sp1, H4Sp2, H5’Sp1, H5Sp1) based on visible differences in the sediment (e.g., colour, texture, and presence of rocks) and the abundance of cultural materials. Horizon 0 represents the topsoil, while Horizon 5 is the last layer excavated.

### 3.2 Studied material

In total, 40 bone artefacts were recovered during the 2021-2024 field seasons. The material was scattered across different archaeological horizons (see Table 1).

**Table 1.** The list of bone artefacts analysed.

| ID | Exc Year | Horizon | Subhorizon | Calibrated Age BCE (95.4%) |
| --- | --- | --- | --- | --- |
| BTL-1 | 2022 | Horizon 0 |  | 3589 – 3528 |
| BTL-2 | 2021 | Horizon 0 |  | 3589 – 3528 |
| BTL-3 | 2021 | Horizon 5 | Subhorizon 1 | 4136 – 4054 |
| BTL-4 | 2022 | horizon 4 | Subhorizon 1 | 3986 – 3914 |
| BTL-5 | 2022 | Horizon 3 | Subhorizon 1 | 3814 – 3704 |
| BTL-6 | 2022 | Horizon 2 | Subhorizon 1 | 3806 – 3704 |
| BTL-7 | 2022 | Horizon 2 | Subhorizon 1 | 3806 – 3704 |
| BTL-8 | 2022 | Horizon 0 |  | 3589 – 3528 |
| BTL-9 | 2022 | Horizon 1 | Subhorizon 1 | 3696 – 3562 |
| BTL-10 | 2022 | Horizon 1 | Subhorizon 2 | 3715 – 3641 |
| BTL-11 | 2023 | Horizon 4 | Subhorizon 2 | 3991 – 3934 |
| BTL-12 | 2022 | Horizon 2 | Subhorizon 2 | 3810 – 3702 |
| BTL-13 | 2022 | Horizon 2 | Subhorizon 2 | 3810 – 3702 |
| BTL-14 | 2022 | Horizon 2 | Subhorizon 2 | 3810 – 3702 |
| BTL-15 | 2022 | Horizon 2 | Subhorizon 1 | 3806 – 3704 |
| BTL-16 | 2022 | Horizon 1 | Subhorizon 1 | 3696 – 3562 |
| BTL-17 | 2022 | Horizon 2 | Subhorizon 1 | 3806 – 3704 |
| BTL-18 | 2022 | Horizon 1 | Subhorizon 1 | 3696 – 3562 |
| BTL-19 | 2022 | Horizon 1 | Subhorizon 2 | 3715 – 3641 |
| BTL-20 | 2022 | Horizon 0 |  | 3645 – 3531 |
| BTL-21 | 2022 | Horizon 0 |  | 3645 – 3531 |
| BTL-22 | 2023 | Horizon 4 | Subhorizon 2 | 3991 – 3934 |
| BTL-23 | 2023 | Horizon 4 | Subhorizon 2 | 3991 – 3934 |
| BTL-24 | 2022 | Horizon 5 | Subhorizon 1 | 4136 – 4054 |
| BTL-25 | 2022 | Horizon 0 |  | 3645 – 3531 |
| BTL-26 | 2022 | Horizon 0 |  | 3645 – 3531 |
| BTL-27 | 2023 | Horizon 2 | Subhorizon 2 | 3810 – 3702 |
| BTL-28 | 2024 | Horizon 5 | Subhorizon 2 | 4142 – 4051 |
| BTL-29 | 2024 | Horizon 4 | Subhorizon 1 | 3986 – 3914 |
| BTL-30 | 2024 | Horizon 2 | Subhorizon 2 | 3810 – 3702 |
| BTL-31 | 2024 | Profile |  |  |
| BTL-32 | 2024 | Horizon 2 | Subhorizon 1 | 3806 – 3704 |
| BTL-33 | 2024 | Horizon 2 | Subhorizon 2 | 3810 – 3702 |
| BTL-34 | 2024 | Horizon 2 | Subhorizon 2 | 3810 – 3702 |
| BTL-35 | 2024 | Horizon 2 | Subhorizon 2 | 3810 – 3702 |
| BTL-36 | 2024 | Horizon 1 | Subhorizon 2 | 3715 – 3641 |
| BTL-37 | 2023 | Horizon 1 | Subhorizon 1 | 4056 – 3971 |
| BTL-38 | 2023 | Horizon 4 | Subhorizon 1 | 3986 – 3914 |
| BTL-39 | 2024 | Horizon 2 | Subhorizon 2 | 3810 – 3702 |
| BTL-40 | 2024 | Horizon 2 | Subhorizon 1 | 3806 – 3704 |

### 3.3 Species identification

All specimens were initially examined morphologically to identify the anatomical element and, where possible, assign them to a specific taxon based on observable morphological traits. In addition, all artefacts were measured using digital callipers. Figure 2 and Supplementary Tables 2 and 3 present the anatomical elements, morphological identifications, and recorded measurements for each artefact.

**Figure 2.**
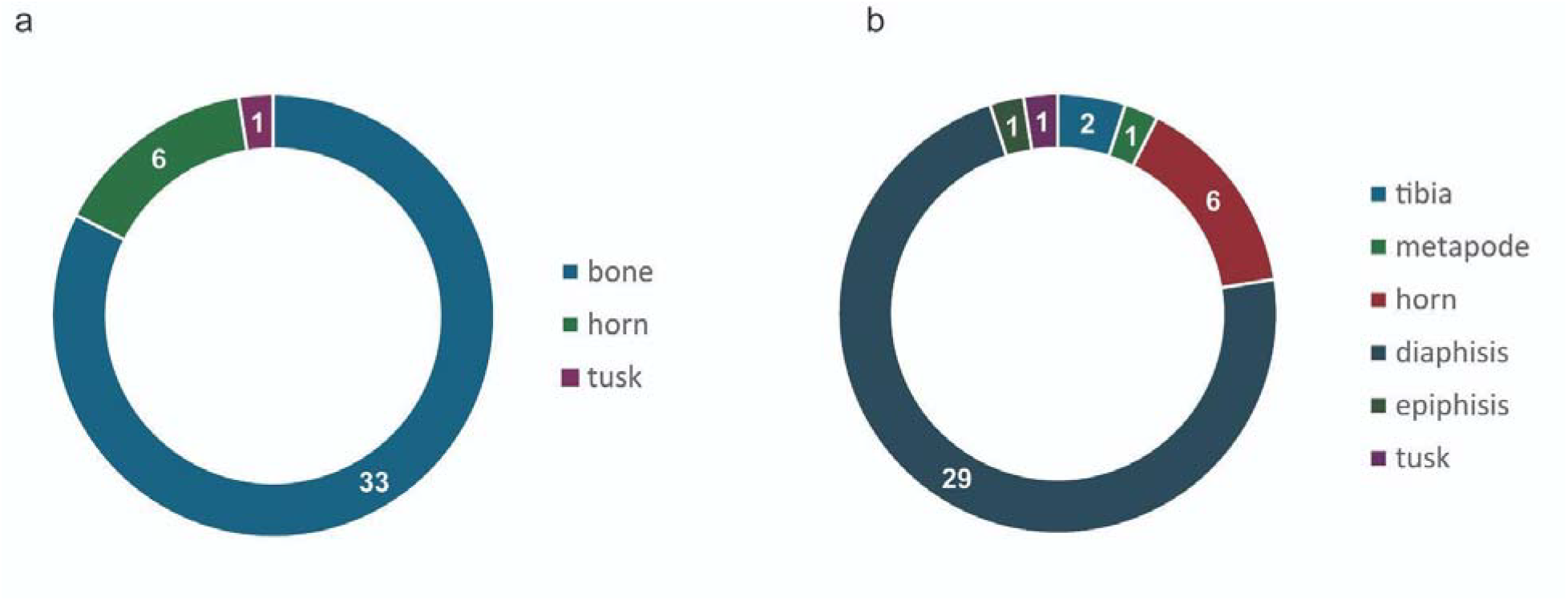
a) raw material choices (n=40); b) anatomical element preferences (n=40)

A minimally invasive sampling technique, using polishing films, was applied to the bone artefacts to avoid visible alterations to the artefact surface. The approach has recently been successfully adapted to artefacts recovered from Neolithic to Iron Age sites across Armenia, demonstrating high identification rates (Antonosyan et al., 2025a). Measures were taken to minimise potential cross-contamination. This included nitrile gloves and face masks, equipment, and workspace cleaning with bleach and ethanol after each sampling. The samples were abraded (1□×□0.5□cm of the bone surface) with coarse polishing film (Fiber Optic Polishing Film by Precision Fiber Products, grit size: 30 μm), causing a small amount of organic material to adhere to the film. Each of these samples was visible as a small amount of powdered material on the surface of the film.

The film containing bone powder was incubated in 100□μ□ 80□mM ammonium bicarbonate solution (AmBic) at 65°C for one hour. Samples were then spun briefly at 9,000□rpm, and 50□μl of supernatant was transferred to a fresh 1.5LmL microcentrifuge tube for subsequent enzyme digestion. The sample was incubated overnight with trypsin solution (0.4□μl/μg), after which 1 μL of 5% Trifluoroacetic acid (TFA) was added to stop enzymatic action. The peptides were purified using C18 ZipTip pipette tips. A total of 3Lμl of collagen extract was mixed with 3□μL of α-cyano-hydroxycinnamic acid matrix. One μl of the sample was spotted in triplicate, each with a different calibrant/reference spot, on a 384-well Bruker MALDI ground steel target plate. The samples were run on a Bruker Autoflex Speed MALDI-TOF mass spectrometer (Bruker Daltonics) to produce spectra/fingerprints for taxonomic identification. ZooMS screening was carried out at the Max Planck Institute of Geoanthropology. The resulting peptide markers were identified via mMass software: v5.5.058. The polishing film displayed low-intensity non-diagnostic background peaks, which did not influence the identifications. The registered collagen fingerprints of each specimen are presented in Supplementary Table 1.

### 3.4 Technological, typological, and use-wear analysis

The technological, typological and use-wear methodologies applied to the study of tools made from hard animal materials, such as mammal bone, teeth and horn, in this research followed the procedures outlined by Poplin (1974), d’Errico and Giacobini (1985), d’Errico (1993), LeMoine (1997), Maigrot (1997, 2003), Averbouh and Provenzano (1999), Averbouh (2000), Cristiani and Alhaique (2005), Sidera and Legrand (2006), Legrand and Sidera (2007) Buc (2011), Alvarez et al., (2014), and Évora (2015). Additionally, we followed the analytical protocols established by the Commission de Nomenclature sur l’Industrie de l’Os Préhistorique (Camps-Fabrer, 1974, 1977), the Fiches Typologiques de l’Industrie Osseuse Préhistorique (Christensen, 2004), and the Multilingual Lexicon of Bone Industries (Averbouh et al., 2015). These references collectively inform the technological, typological, and use-wear analysis approaches as well as the identification of different categories of artefacts (such as blocks, blanks, preforms, and finished objects), and the application of the *chaîne opératoire* approach (Averbouh 2000), providing the reconstruction of the reduction sequence.

All artefacts were initially examined macroscopically and subsequently analysed under a binocular stereomicroscope with magnification levels ranging from 10x to 40x. Photographic documentation was also carried out for each specimen, and measurements were taken using a digital calliper. In this paper, all measurements are presented in millimetres (mm). Key variables recorded include cortical thickness, presence or absence of spongy tissue, fracturing marks and techniques, fracture planes, and surface modifications or use-wear traces. The manufacturing techniques, morphological variability, and use-wear patterns observed on the bone tools allow us to infer their possible functions. These aspects, taken together, contribute to a better understanding of raw material selection and the functional purposes of osseous artefacts used by the Chalcolithic communities in this region of Armenia. A basic taphonomic analysis was also conducted to verify natural surface alterations, following the guidance of Behrensmeyer (1978), Behrensmeyer et al., (1986), Lyman (1994), Fisher (1995), Blumenschine et al., (1996), Dominguez-Rodrigo et al., (2012), and Fernandez-Jalvo and Andrews (2016). This task was essential for distinguishing intentionally modified tools from pseudo-tools (Binford, 1981) (objects that bear use-like traces caused by non-cultural processes). These pseudo-tools typically exhibit trampling marks and randomly distributed polish or abrasion resulting from contact with taphonomic agents such as water, sediments, wind erosion, and carnivore action.

## 4 Results

### 4.1 Taxonomic composition

A total of 40 bone artefacts were analysed in this study. All of the specimens were first morphologically screened to identify the anatomical element and, where possible, identified to a taxon based on anatomical characteristics. In most cases, extensive modifications resulting from artefact production limited taxonomic identification, and samples were therefore classified into broad size categories (e.g., large mammal, small mammal). Only in a few instances it was possible to assign specimens to a genus level based on morphology. These included sheep and goat horn fragments (BTL-1, 5, 18, and 28), a tibia identified as Lepus (BTL-4), a tine attributed to either deer or antelope (BTL-21), and a tusk of a boar (BTL-29). One specimen (BTL-3) was identified as a Caprine metapodium.

The majority of the artefacts were shaped from the long bone diaphyses of large and small mammals (n=29; Figure 2). Other anatomical elements represented include epiphysis (n=1), horns (n=4), metapodium (n=1), tibia (n=2), tine (n=1), and tusk (n=1).

ZooMS was conducted on all 40 bone artefacts using a minimally invasive sampling, of which 39 preserved enough collagen to allow for taxonomic identification, while only one failed to yield sufficient collagen for analysis. In the majority of cases, identifications were made at the genus level (n=36). In a few instances (n=3), limited preservation constrained identifications to the family level. For instance, three specimens (BTL-8,10 and 11) could only be classified as “Bovidae” through ZooMS due to poor preservation, resulting in a lack of diagnostic markers.

In total, six taxonomic groups were identified, aligning well with the zooarchaeological findings from the site (Antonosyan et al., 2025b). The results of the taxonomic identification through ZooMS are represented in Figure 3, and the recorded ZooMS markers are available in Supplementary Table 1. The identified assemblage is dominated by Caprines: 16 of the analysed tools were manufactured from goat (*Capra sp.*) bones and 11 from sheep (*Ovis sp.*) bones. ZooMS enabled this distinction between sheep and goat through the COL1α2 757–789 marker, identified at m/z 3014.4 and 3033.4 for sheep, and m/z 3077.4 and 3093.4 for goats.

**Figure 3.**
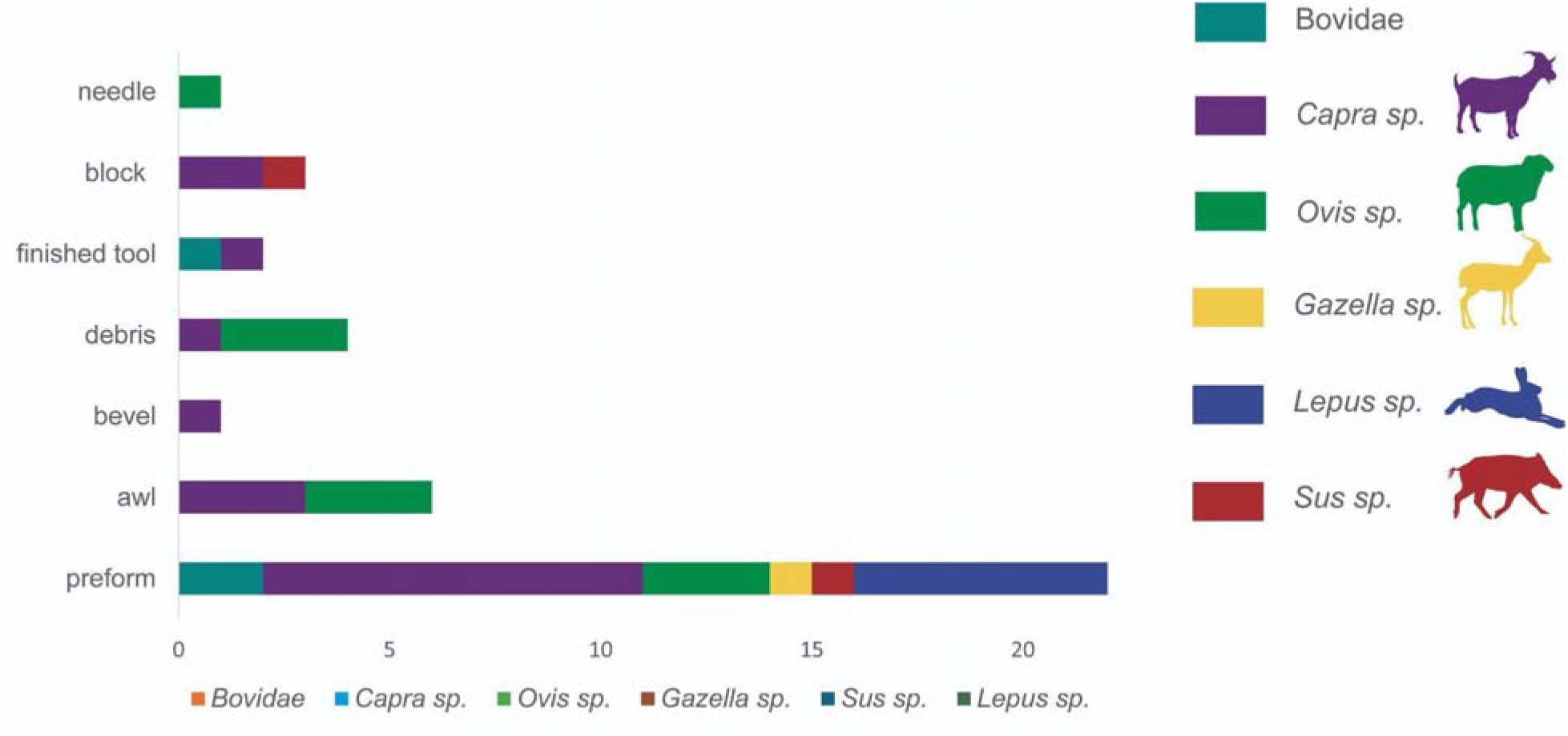
Taxa used in artefact manufacture

ZooMS also proved effective in distinguishing members of the Antilopini tribe, using the COL1A2 375 and α2 889 markers, which for Antilopini are characterised by m/z values of 1182, 2056, 2072, and 1532, respectively. Based on these markers, one artefact (BTL-21) was identified as being made from Gazella bone, providing clear evidence for the use of wild game in bone tool production. Additionally, two artefacts (BTL-39 and BTL-29) were made from boar bone (*Sus sp*.). Another frequently represented group is hare (*Lepus sp.),* which was used in the manufacture of six artefacts.

ZooMS analysis provided insights into raw material selection across different artefact types. Goat bones were used both as a block and for the production of finished tools, such as awls, and were also identified among tube preforms, bevels, and manufacturing debris. Sheep bones were selected for the production of tools like needles and awls and were similarly represented in the form of tube preforms and debris. Boar bone appeared as both a block and a preform. A single gazelle specimen was identified as a preform, as were all hare bones.

The distribution of identified taxa across the six horizons is shown in Figure 4. Sheep, goats, and hares are present consistently throughout the sequence. Gazelle appears only in the uppermost layer, while boar seems to emerge in the later phases of the sequence.

**Figure 4.**
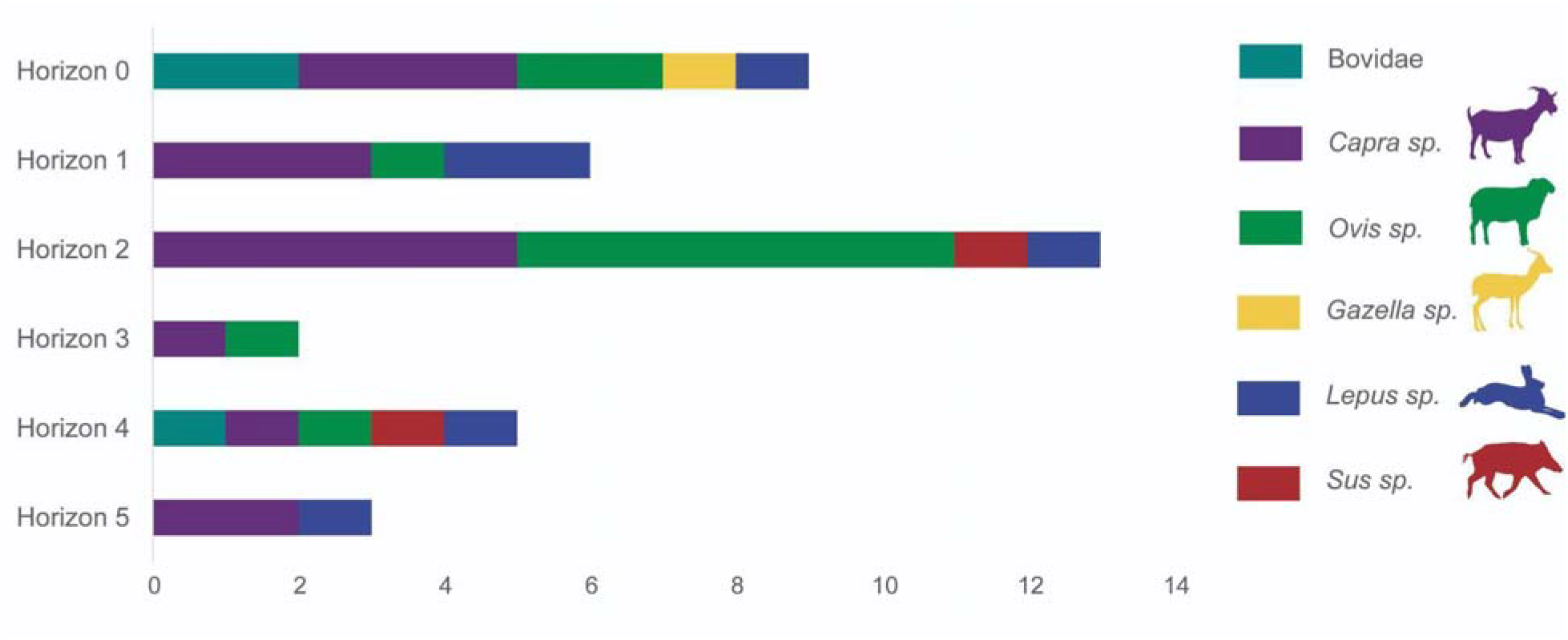
Distribution of identified taxa across the six horizons.

### 4.2 Technological analysis

The technological analysis enables the identification of fracture patterns and, consequently, provides insights into raw material procurement strategies and economy. In the assemblage analysed in this study (n=40), several osseous artefacts exhibit varying degrees of technological transformation. The data shown in Supplementary Tables 2 and 3 include variables such as animal species of origin, anatomical element choices, tool categories, condition of the artefact, cortical tissue thickness, and the transformation techniques applied. Following this data, we have identified some morphological and technological patterns to understand the technical and functional choices behind the manufacturing process of these tools.

The presence of the full *chaîne operatoire* is demonstrated by the identification of categories such as block, preform, debris, and finished tools, and the fracturation and modification techniques identified include sawing, bending, percussion (direct and indirect), and abrasion, with some combinations between them (Figure 5). The most frequently employed transformation technique was sawing the bone diaphysis, followed by bending the extremities, applied predominantly to obtain tube preforms, indicating an initial stage of the transformation sequence. Other techniques, such as percussion and abrasion, were associated with thicker pieces, such as awls, a bevel, and a smoother.

**Figure 5.**
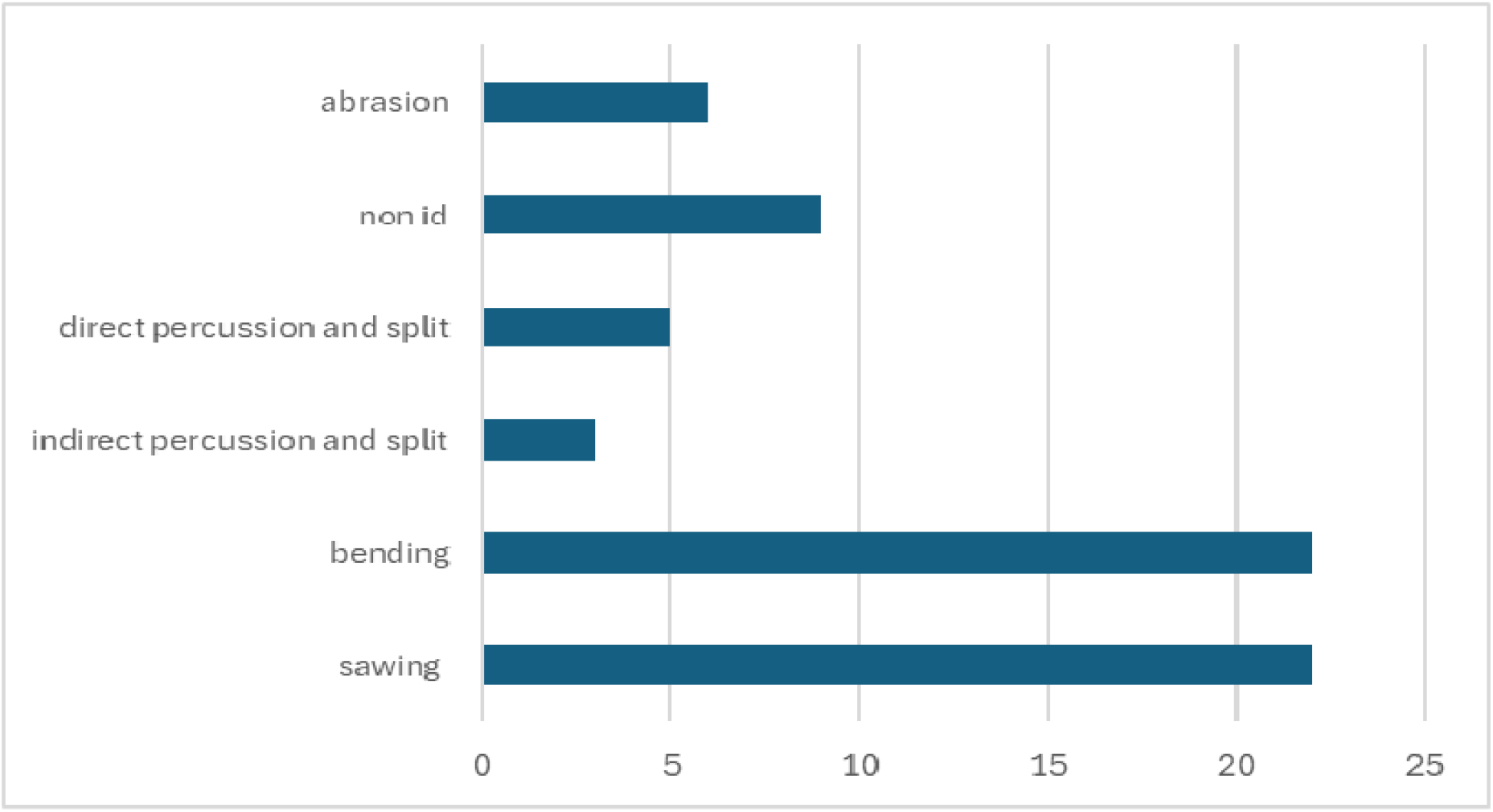
Fracturation and modification techniques applied to raw materials (non-id = not non-identified).

The analysis of the cortical thickness enabled an indirect assessment of bone density and its relationship to the type of transformation technique applied. The predominance of anatomical elements such as the diaphysis, with an average cortical thickness of around 3.2 mm, may reflect a deliberate choice for long and thick bones suitable for prolonged use of the tools obtained, as is the case with a bevel and some of the awls. Meanwhile, bones with thinner cortical tissues were suitable for sawing and bending to obtain the desired preform. This association between anatomical elements and transformation techniques points to a sophisticated empirical knowledge of the available bone materials, their mechanical properties, and their technological potential.

### 4.3 Use-wear analysis

Through the use-wear analysis, we verified the existence of different marks on the bone surface of the utensils, some resulting from the manufacturing techniques applied and others due to the probable function(s) of the artefact (Figure 6). Most of the tools still have abrasion marks on their surface, and these seem to result from using an abrasive sediment or stone of moderate coarseness.

**Figure 6.**
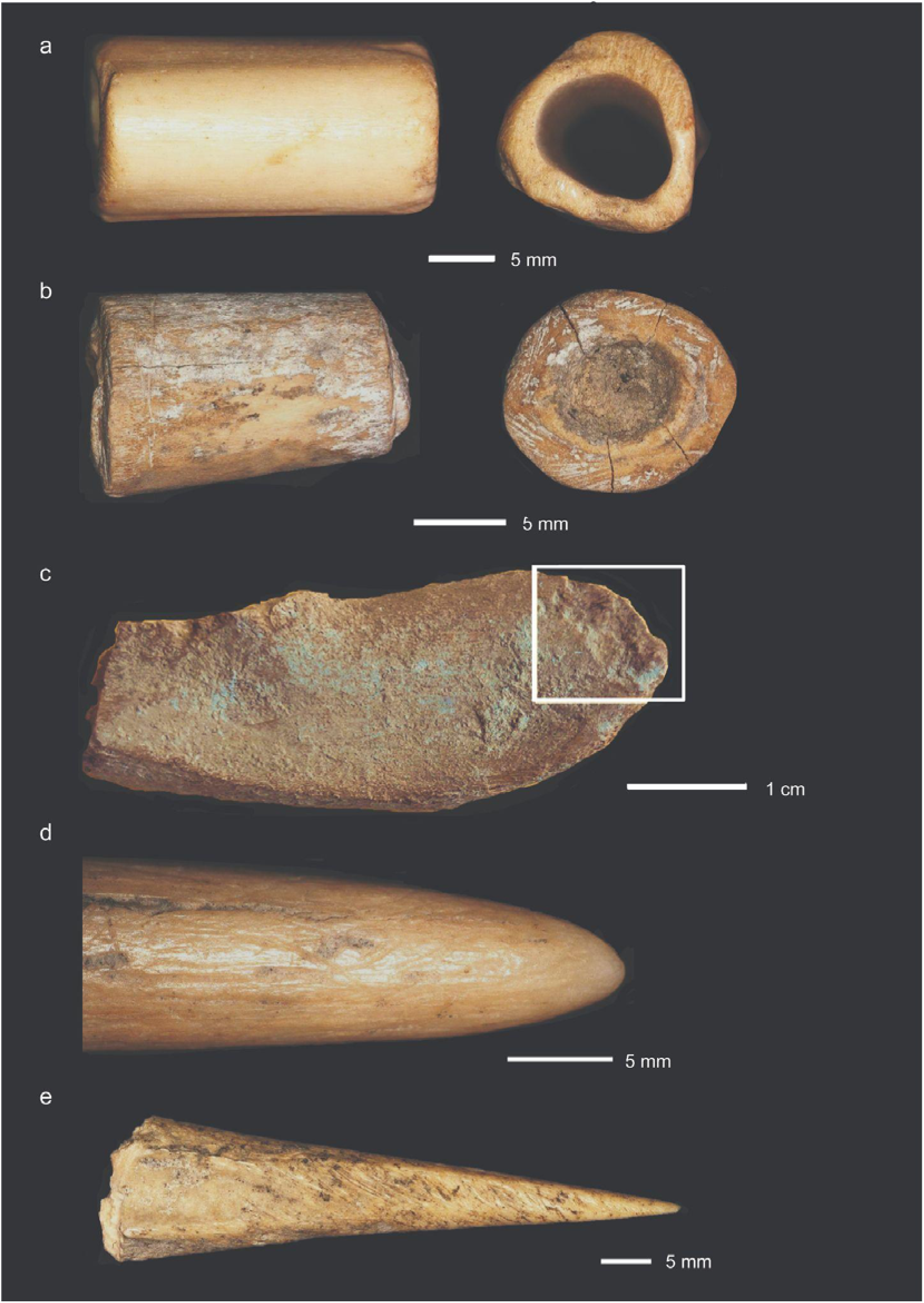
: a) Tube preform made from Lepus sp. tibia showing sawing marks (on the right-hand side). b) Tube preform made from Gazelle sp. horn showing sawing marks (on the left-hand side) and bending marks (on the right-hand side). c) Bevel made from Capra sp. diaphysis, its active area (marked in the photo) shows the negatives of bone removals resulting from impacts. d) Needle tip made from *Ovis sp.* diaphysis with use-wear *stria* on its active area. e) Fragment of an awl made from Capra sp. diaphysis with use wear marks on the bone surface.

The tube preforms have sawing striae on their extremities, and some also have abrasion striae on their extremities that removed the bending marks. No other modification techniques were applied on their surface, except for one tube preform whose both extremities were abraded after the sawing was performed. Its outer surface has several small incisions with different orientations, some aligned in a vertical line, others in a diagonal line, transverse to the long axis of the bone tube. None of the tube artefacts are finished.

The smoother artefact has slight grooving marks on the top of the bone surface, with a diagonal direction, and some of them have the Λ orientation (representing a go/stop/return movement). Its active area occupies the whole surface of the bone fragment. The retoucher contains an active area with short incisions all around the proximal end. On its distal end, which is more or less pointed, has negatives from impact marks (small removals of the bone surface). The proximal end of this artefact was possibly used as a retoucher. Regarding the needle, the tip is intact. There are abrasion *striae* on the fragment bone surface with a random orientation, and use-wear marks on top of these *striae* have a horizontal and diagonal orientation, regarding to the long axis of the bone needle, and are concentrated in an active area length of about 25mm from the tip of the needle to the middle of the fragment.

On the awls, one of the extremities of the bone artefacts was abraded to form a point later used as a perforation tool; all have *striae* on the top of the bone surface with longitudinal and vertical orientation parallel and transversal, respectively, to the long axis of the bone. Some awls have small *striae* with random orientations in addition. Others have their tip broken, probably during their use. One awl contains an active area with short incisions all around the proximal end and also impact marks (negatives from impacts, small removals of the bone surface), and its distal end is more or less pointed; this artefact, in particular, was possibly used as a retoucher on the proximal end, and as an awl in the distal end. These awls could have been used to work on leather and textile, and as manipulators in the making of basketry (Campana, 1989).

The bevel has small negative removals of bone in the distal active area due to its function, and they are present on the outer and inner surfaces of the bone fragment. This tool could have been used for woodworking and/or bone working. The tusk block is a proximal-mesial fragment, still with the brownish colour on one of its extremities (proximal). It was split into two longitudinal parts, and on its lateral sides, there are abrasion striae with a diagonal orientation. Its extremities do not present marks of use-wear or modification, which could be a discard.

Regarding the basic taphonomic analysis of the bone artefacts in this assemblage, we could confirm that none of the artefacts analysed was a pseudotool, although we registered the presence of root marks, manganese dioxide staining, burnt surfaces, longitudinal fractures, small tooth marks, random orientation *striae* on some parts of the bone surfaces in a few tools but not in the active areas, and sediment concretions.

## 5. Discussion

As this is a preliminary study and given the available sample size per horizon, we cannot yet safely indicate preferences for raw materials selection over time. The strategies for animal species selection to manufacture these tools are probably influenced by the properties of the skeletal elements from the available fauna around the site. Sheep, goats, and hares are present consistently throughout the sequence. *Gazelle sp*. appears only in the uppermost layer, while *Sus sp.* seems to emerge in the later phases of the sequence.

The most exploited species for bone tool manufacture was *Capra sp*., followed by *Ovis sp*., both representative of pastoral and agricultural contexts. While ZooMS is effective in distinguishing these closely related species, it does not allow differentiation between wild and domestic forms. During the Chalcolithic, both sheep and goats were fully domesticated, and stable isotope data from the site suggest that these animals were managed by herders (Antonosyan et al., 2025b). However, wild relatives of both species were also present in the region, making it difficult to determine with certainty whether the identified Caprines represent domestic or wild taxa. Similarly, it remains uncertain whether the *Sus sp.* tusk derives from domesticated pigs or wild boars. Although domestic pigs have been reported at the contemporaneous site of Godedzor (Palumbi et al., 2021), the timeline for pig domestication in the region is still under debate. This restricts the understanding of strategies in selecting raw materials for bone tool production for these species.

In this assemblage, diaphysis was the main anatomical element of choice to produce bone tools carved on ungulate bones, such as tube preforms, awls, bevels, and needles. In contrast, *Lepus sp*., displaying a much lower cortical tissue thickness, may have limited its use to lighter or maybe more specific tools, although the artefacts analysed in this assemblage are also preforms but not as thick as preforms made from *Capra sp*. or *Ovis sp.* diaphyses.

The versatility of horn is evidenced by a wide thickness range (from 0.66 to 9.42 mm), suggesting it was exploited due to its flexibility. Horns were utilised as a block, a preform, and debris, as indicated by their *chaîne operatoire* at this site. A *Gazelle sp*. preform made from horn with 4.95 mm cortical thickness was sawn in the middle of the tine, to obtain a tube. The tubes are the most relevant type of preform in this assemblage, although they are not perforated, and are not finished either. This, along with the high frequency of this tool, shows possible evidence of manufacturing specialisation at the site, and this production could be directed to personal use among the inhabitants around Yehegis-1 or even for exchange networks. Their function is still debatable, and Averbouh (1993) indicates 3 possible functions for this type of tool: as a musical instrument, as a recipient, and as an amulet or personal ornament.

The prevalence of fracturation and wear techniques such as bending and sawing on the diaphysis to obtain the preforms of tubes, and the association of techniques like percussion on thicker cortical tissue bones, all reflect a well-defined technological strategy. Thicker cortical tissue bones were selected for procedures that involved higher structural stress during their use, like the bevel and some awls. The relationship between cortical tissue thickness, anatomical element type, and the transformation and modification techniques indicates a technically informed practice, based on the functionality and durability of the final artefacts. The association between anatomical elements and the applied fracturation technique was important to understand the technological knowledge involved in the manufacturing process. The osseous tools *chaîne opératoire* present at the Yeghegis-1 rock shelter indicate that people were producing the tools they needed for their daily tasks at the site, such as needles, awls, bevels, and smoothers, from the gathering of the blocks to the finished object.

Based on this preliminary data from Yeghegis-1 rock shelter, the bone tool assemblage reflects typological and technological patterns characteristic of Chalcolithic communities in the territory of Armenia, with parallels in sites such as Areni-1 Cave, Getahovit-2 Cave, Aknashen-Khatunarkh and Godedzor sites, and more distant regions, including Godin Tepe (Iran) (Crabtree and Campana, 2010), Uğurlu and Tepecik-Çiftlik (Anatolia) (Kalantaryan and Ghanem, 2019; Paul and Erdoğu, 2017), Polyanitsa tell (Northeastern Bulgaria) (Shakun et al., 2025), VităneLti Măgurice (Romania) (Mărgărit, et al., 2022), and Yiftahel Area C (Southern Levant) (Garfinkel et al., 2012). The recurrence of standardised forms, like awls, bevelled tools, and tubes, suggests not only functional convergence but also possible cultural transmission mechanisms and shared craft traditions across cultural boundaries. This standardisation may also be indicative of interregional exchange networks active during the Chalcolithic, through which ideas and finished tools (and eventually raw materials) circulated.

## 6. Conclusions

This preliminary study of the osseous tool assemblage from Yeghegis-1 provides detailed insights into bone tool production strategies and species selection for bone manufacturing in Chalcolithic Armenia. The results reflect a possible preference for some osseous raw materials, their transformation, and modification techniques. The use of a minimally invasive sampling technique for ZooMS screening proved effective, as it enabled the identification of animal species while preserving the integrity of cultural heritage objects. The ZooMS technique allowed the taxonomic identification of the animal species from which the utensils were manufactured, as being dominated by sheep and goats. These results are in agreement with the previously established zooarchaeological estimates at the site, which suggested that animal husbandry, particularly of sheep and goats, was an important component of the subsistence economy. The presence of bones of wild game, such as hare and gazelle, at the site points to a reliance on hunted taxa for the production of artefacts as well. Gazelles and hares have been identified in Yeghegis-1 through the ZooMS screening of faunal assemblage, though in smaller numbers in comparison to the domesticated fauna. This suggests that the early pastoralists in the region mostly utilised readily available resources for bone manufacture.

Our analysis also contributes to broader discussions regarding resource management and its application represents an important tool for understanding the socio-economic dynamics of prehistoric communities. The technical patterns identified in this preliminary study reflect an understanding of the available raw materials, as well as an operational logic that includes technique, form, and function. These observed patterns support the hypothesis of the existence of a small-scale workshop operating at the site, corroborated by an intentional and specialised *chaîne opératoire* in osseous raw material transformation, consistent with systematic production practices of diverse tools used in the daily activities of these communities. Further, similarities with assemblages from Armenia and surrounding regions can point to the circulation of knowledge and the existence of shared craft traditions, sustained through interregional exchange networks operating during the Chalcolithic. Such networks likely played a critical role in shaping shared technological traditions and fostering contacts among communities across a broad geographical area.

## Acknowledgments

We are grateful to the staff of Vayots Dzor Regional Museum in Yeghegnadzor, Armenia, whose generous support has made the work possible. We thank Karine Stepanyan, the director of Yeghegnadzor Regional Museum. We are grateful to Dr. Yoshi Maezumi (Max Planck Institute of Geoanthropology) for providing access to the binocular stereomicroscope used in this study.

## Author Contributions statement

**M.A.:** project administration, conceptualisation, investigation, data curation, formal analysis, visualisation, writing original draft, review and editing; **M.E.:** conceptualisation, data curation, formal analysis, investigation, writing original draft, review and editing; **S.M.:** formal analysis, review and editing; **M.S.:** review and editing; **N.A.:** review and editing; **U.T.:** visualisation; **S.Sh:** review and editing; **L.Y.:** review and editing; **P.R.**: resources, review and editing.

